# A Node-based Informed Modularity Strategy to Identify Organizational Modules in Anatomical Networks

**DOI:** 10.1101/2020.07.06.189175

**Authors:** Borja Esteve-Altava

**Author notes:** Barcelona Biomedical Research Park, Doctor Aigüader 88, 08003 Barcelona, Spain. /.

## Abstract

The use of anatomical networks to study the modular organization of morphological systems and their evolution is growing in recent years. A common strategy to find the best partition of anatomical networks into modules is to use a community detection algorithm that tries to optimize the modularity Q function. However, this strategy overlooks the fact that Q has a resolution limit for small modules, which is often the case in anatomical networks. This produces two problems. One is that some algorithms find inexplicable different modules when we input slightly different networks. The other is that algorithms find asymmetric modules in otherwise symmetric networks. These problems have discouraged researchers to use anatomical network analysis and boost criticisms to this methodology. Here, I propose a *Node-based Informed Modularity Strategy* (NIMS) to identify modules in anatomical networks that bypass resolution and sensitivity limitations by using a bottom-up approach. Starting with the local modularity around every individual node, NIMS returns the modular organization of the network by merging non-redundant modules and assessing their intersection statistically using combinatorial theory. Instead of acting as a black box, NIMS allows researcher to make informed decisions about whether to merge non-redundant modules. NIMS returns network modules that are robust to minor variation and does not require to optimize a global modularity function. NIMS may prove useful to identify modules also in small ecological and social networks.

## INTRODUCTION

Anatomical network analysis has recently emerged as a new framework to study anatomy quantitatively using tools from network theory (Rasskin-Gutman & Esteve-Altava, 2014). This approach first formalizes anatomical systems as network models, in which nodes represent individual anatomical elements (e.g., bones) and links represent pair-wise relations among them (e.g., articulations), and then quantifies their topological organization as a proxy to understand the biological features of the body. Anatomical studies using network analysis have focused on comparing the development, function, and evolution of morphological systems, from invertebrates to vertebrates, including extant and extinct organisms (e.g., Esteve-Altava, Marugán-Lobón, Botella, & Rasskin-Gutman, 2013; Esteve-Altava, Molnar, Johnston, Hutchinson, & Diogo, 2018; Esteve-Altava et al., 2019; Saucède et al., 2015; Dos Santos, Fratani, Ponssa, & Abdala, 2017; Kerkman, Daffertshofer, Gollo, Breakspear, & Boonstra, 2018; Murphy et al., 2018; Ostachuk, 2019; Plateau & Foth, 2020).

Modularity is one of the most explored features of anatomical networks. In this context, a module is a group of nodes with more connections among them than to other nodes outside the group. We can classify network-based modules as organizational modules: *“Organizational morphological modules refer explicitly to the interactions postulated to be important in organismal construction or activity. They invite observation or description in terms of mechanistic relations, whether variation among organisms is present or not. As such, organizational modules are units of stability”* (Eble, 2005). The vertebrate skull is the most frequently object of study when looking for morphological modules (Esteve-Altava, 2017b). In this context, anatomical network studies have tried to unveil the topological units of organization to understand how they vary in evolution (Esteve-Altava, Boughner, Diogo, Villmoare, & Rasskin-Gutman, 2015; Arnold, Esteve-Altava, & Fischer, 2017; Werneburg, Esteve-Altava, Bruno, Torres Ladeira, & Diogo, 2019; Plateau & Foth, 2020), but also in normal and pathological development (Diogo et al., 2019). Moreover, some other studies have applied network methods to identify modules of landmark-based morphometric correlations rather than topological relations (Ivan Perez, de Aguiar, Guimarães, & dos Reis, 2009; Suzuki, 2013). A connection between network-based organizational modules and shape covariation modules would be likely (Esteve-Altava, Marugán-Lobón, Botella, Bastir, & Rasskin-Gutman, 2013), but it has not been elucidated yet (Esteve-Altava, 2017a).

A common strategy to delimit modules in anatomical networks is to use a community detection algorithm that optimize a quality function, which measures how well divided is the network into modules compared to a random model. The most popular optimization function is the modularity Q (Newman & Girvan, 2004). Q measures the number of links within modules compared with a random distribution of links between all nodes regardless of modules. However, Q has a resolution limit for smaller modules that depends on the size of the network and the possibility to distant nodes to connect (Fortunato & Barthelemy, 2007). As a result, optimization algorithms may fail to identify small modules in relatively larger networks. This is because in large networks connections cannot be distributed purely at random. Although anatomical networks are smaller compared to other natural networks (e.g., genetic, ecological), the resolution limit applies here because geometry and development constrain connectivity (Esteve-Altava & Rasskin-Gutman, 2014). Despite their limitations, optimization algorithms are persistently used to study anatomical networks because they are easy to apply and because they are readily available through build-in packages and programs like *igraph* (Csardi & Nepusz, 2006). Other resolution problems originate from the fact that many nodes have spread their links evenly between modules. Because most nodes tend to have a small number of connections, this makes difficult to delimit modules solely by connections in and out of modules.

Global optimization and resolution limits produce two problems in many modularity studies of anatomical networks. The first problem is that algorithms “inexplicably” find different modules when we input slightly different networks. This happens, for example, when one fixes a minor mistake in a network and finds out that in the revised network the modularity output changes considerably. It also happens when comparing similar networks with intraspecific variation (e.g., adding a Wormian bone in a skull), for which we would expect no major changes of modularity. The second problem is that algorithms sometimes find asymmetric modules in otherwise symmetric networks, which challenges basic biological assumptions about symmetry of bilateral body parts. These problems can hinder anatomical network analyses and they fuel criticisms to this methodology from people unaware of these limitations.

Here I propose a strategy to bypass resolution and sensitivity limitations, using a bottom-up approach that frames the problem of finding modules in anatomical networks at the level of individual nodes: a *Node-based Informed Modularity Strategy* (NIMS). Starting with the local modularity around every single node, NIMS returns the modular organization of the entire network by merging non-redundant modules and statistically assessing their degree of intersection. Instead of acting as a black box, NIMS allows to make informed decisions about whether to merge non-redundant modules or keep them as separated modules, based on the significance and the size of the intersection. NIMS returns network modules that are robust to minor variations and avoids common pitfalls of other approaches by not requiring to optimize a global modularity function. The following section describes NIMS in more detail. Then, the modularity of four skull networks is analyzed as case studies: (1) human, (2) human with intraspecific variation, (3) tinamou, and (4) crocodile.

## NODE-BASED INFORMED MODULARITY STRATEGY (NIMS)

NIMS builds upon existing functions implemented in R packages (R Core Team, 2019). This has been done for convenience because R is a popular programming language for anatomical network analysis and among biologists. However, NIMS is independent of the programming language we used and other implementations are possible.

NIMS has six sequential steps:

1. Get all node-level modules
2. Remove extra node-level modules with same elements
3. Remove node-level modules with all elements included in a larger module
4. Test multiple-set overlaps
5. Merge node-level modules (top p-value / max overlap size)
6. Test cohesion for network-level modules (Wilcoxon test)

Step 1 delimits node-level modules for every node in the network. This can be done using the function *cluster_spinglass* in the package *igraph* (Csardi & Nepusz, 2006), setting the argument vertex sequentially to every node of the network. This function finds node-level modules based on their difference between realized and expected internal links (cohesion) and between realized and expected external links (adhesion) (Reichardt & Bornholdt, 2006). Alternatively, we could use any algorithm that returns node-level modules, even those based on criteria.

Steps 2 and 3 cross-check node-level modules to filter out modules with the same nodes (duplicated) and modules entirely included as part of a larger module (nested). This produces a list of non-redundant node-level modules. Finding only one module after Steps 2 and 3 means that the nodes of the network are fully integrated and that the network has no modules.

Step 4 evaluates the intersection of non-redundant node-level modules to reveal potential merges of highly overlapping modules. This procedure is co-opted from research on gene sets functional enrichment analyses (Subramanian et al., 2005), extended to perform multiple-set comparisons (for details, see Wang, Zhao, & Zhang, 2015). Step 4 returns the size of the overlap (how many nodes) for all combinations of node-level modules and the statistical significance of their overlap compared to random expectations. In R, we can use the function *supertest* in the package *SuperExactTest* (Wang et al., 2015), which performs multi-set enrichment analysis and provides tools for efficiently plotting the results so we can inspect them visually. To determine when a combination of modules is significant, we must first account for multiple testing using an appropriate p-value correction, such as the Bonferroni’s correction (Miller, 1966). Any combination of modules above the Bonferroni corrected p-value is well supported statistically. P-value thresholds can be incorporated in the visualization of Step 4 to facilitate the visualization of supported merges.

Step 5 requires the researcher to make an informed decision on whether to merge two or more overlapping modules, based on the summary statistics from Step 4. As a rule, we can merge modules into a single module when the overlap is large compared to the number of nodes of each node-level module and when the merging is statistically well supported. Similar well supported merges are possible. For these cases, researchers will have to decide and justify the rationale of the decision.

Finally, Step 6 will test whether the final modules fulfill the definition of network module as a group of nodes with more links in that out the module. We can do that with a paired Wilcoxon signed rank test of the null hypothesis that in-module links are equal than the out-module links (not a module), against the alternative hypothesis that in-module links are greater than out-module links (a module). This test is implemented in the function *wilcox.test* of the *stats* package (R Core Team, 2019).

The next sections demonstrate the application of NIMS to resolve the modular organization of four different anatomical networks. **Supplementary File 1** includes the data and R code to reproduce these examples, as well as an example using the benchmark social network Zachary’s karate club (Newman & Girvan, 2004).

## CASE STUDIES

Four anatomical networks were analyzed representing (1) the ‘type’ adult human skull, (2) a human skull with intraspecific variation, (3) the adult skull of a tinamou, and (4) the adult skull of a crocodile (**Figure 1**).

**Figure 1.**
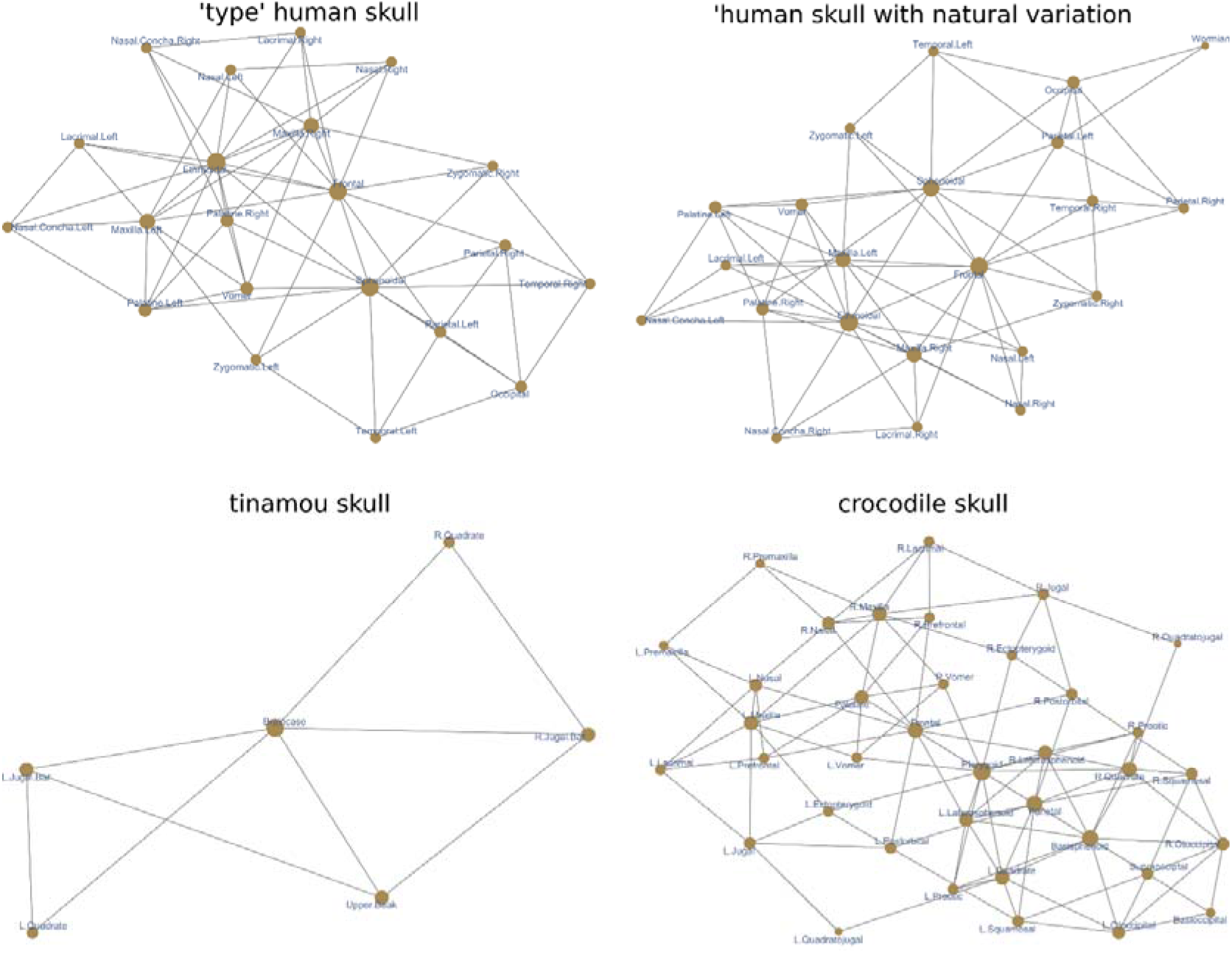
Anatomical networks used as examples.

The first network is the ‘type’ adult human skull and it has been analyzed in previous works (Esteve-Altava et al., 2013; Esteve-Altava et al., 2015). Different methods have consistently identified two modules: one posterior, grouping the bones of the cranial vault and base; and one anterior, grouping the bones of the facial region. Using a local optimization method (Lancichinetti, Radicchi, Ramasco, & Fortunato, 2011), which can find overlapping modules (covers), another study found that these two modules overlap in the frontal and zygomatic bones (Esteve-Altava, 2017a). However, results of this algorithm were sensitive to changes in internal arguments. The analysis of the ‘type’ human skull will serve to compare NIMS to the modules delimited by other algorithms. The expected output is that NIMS can identify the same two modules.

The second network is a modification of the first one that includes anatomical variation found in natural populations (Berry & Berry, 1967). It has an extra Wormian bone between the left parietal and the occipital. It also has different configurations of the pterion region on each side: on the left, the parietal contacts with the sphenoid which prevents the contact between temporal and frontal; on the right, the temporal contacts with the frontal which prevents the contact between parietal and sphenoid. As a result, the network is asymmetric along the left-right axis for the number of links and bones. This network is also an example of normal variation that we find when studying actual skulls rather than type forms. The analysis of this network will show how NIMS deals with intraspecific variation naturally present in anatomical systems, and whether the resulting modules are equivalent to those of the ‘type’ skull network. The expected output is that NIMS has a robust behavior for small variations and finds a modular organization that is consistent with that of the type network.

The third network is the skull of an adult tinamou *Nothura maculosa*, which consist of only six bones due to a process of fusion during postnatal development. Because of the small size of this network, preliminary reports (Lee, Esteve-Altava, & Abzhanov, 2020) found either that this network is not modular (i.e., they find one module) or a trivial partition of the network. The analysis of this network will show how NIMS deals with small, non-modular networks. The expected output is that NIMS find one module.

The fourth network is the adult skull of the crocodile *Crocodylus moreletii*, which consist of a skull with many unpaired bones typical of non-avian archosaurs. For anatomical networks with many bilaterally symmetric nodes, community detection algorithms by optimization sometimes return oddly asymmetric modules (e.g., a module grouping only right-side nodes plus one left bone) or trivially asymmetric by placing unpaired nodes with a one-side module when it is equally connected to the left and right sides (Lee et al., 2020; Plateau & Foth, 2020). The analysis of this network will show how NIMS deals with larger networks with many unpaired bones. The expected output is that NIMS will return a modular organization without asymmetries.

## RESULTS & DISCUSSION

### Modularity of the ‘type’ Human Skull Network

NIMS returns four non-redundant node-level modules, labeled accordingly as occipital, sphenoidal, frontal, and ethmoidal. Note that the names of the node-level modules are taken from the first node which local module is non-redundant and it is merely a label. The statistical analysis of the intersections of the four modules returned two significant overlaps (**Figure 2**), which supports merging node-level modules Ethmoidal + Frontal (labeled facial) and Sphenoidal + Occipital (labeled cranial). The two modules overlap in the frontal, palatines, and vomer; therefore, we can define them as covers (i.e., overlapping modules). The facial cover groups the ethmoid, frontal, lacrimals, maxillas, inferior nasal conchae, nasals, palatines, and vomer. The cranial cover groups the frontal, occipital, palatines, parietals, sphenoid, temporals, vomer, and zygomatics. Statistical evaluation of the two covers confirms they meet the standard definition of module, as a group of nodes with more links in than out the module (facial cover: *W* = 161.5, *p-value* = 3.12e-05; cranial cover: *W* = 125.5, *p-value* = 8.38e-04).

**Figure 2.**
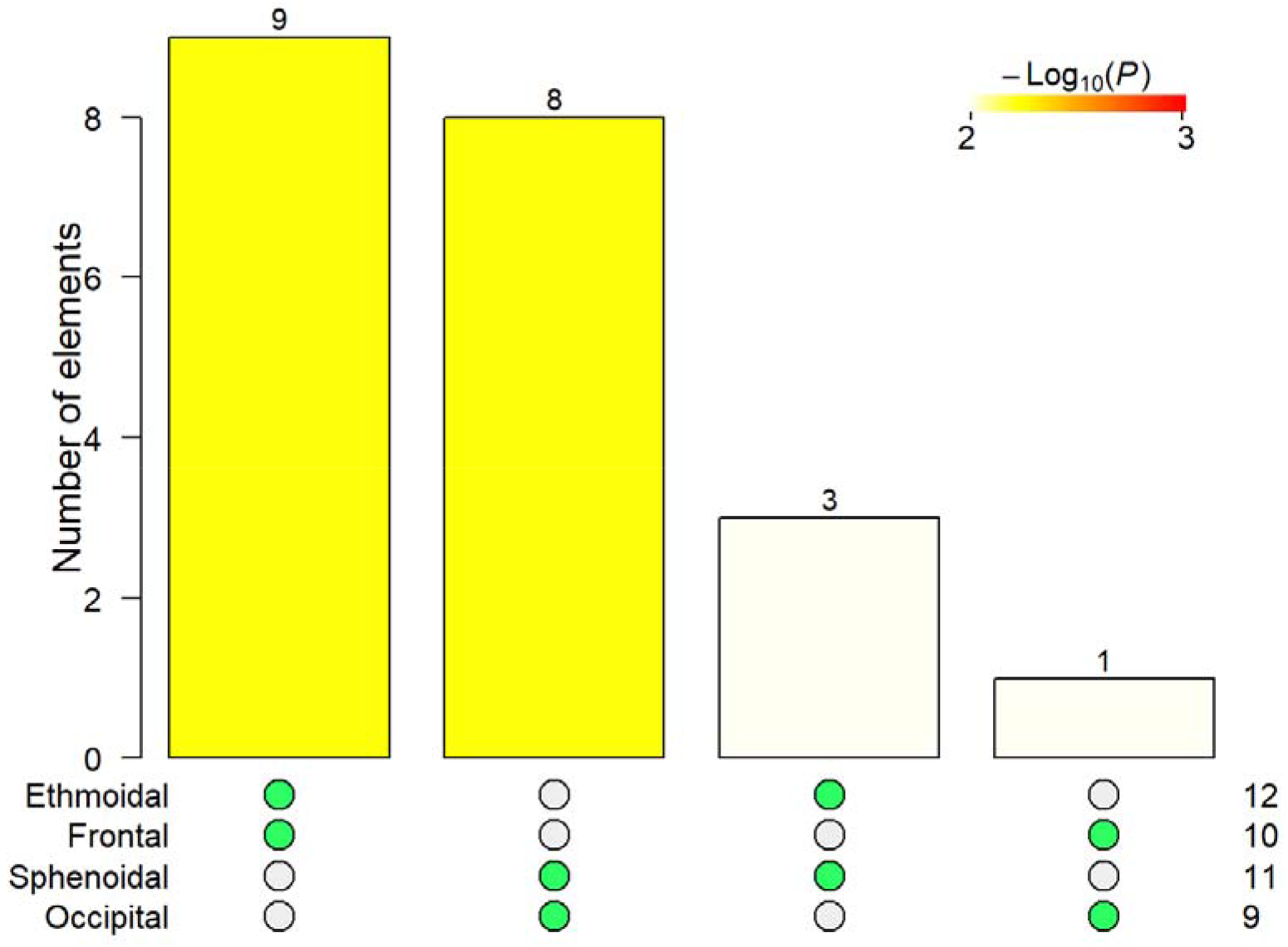
Intersection tests for the four non-redundant node-level modules of the ‘type’ adult human skull network. Green dots indicate the intersection of modules tested and the high of the bar shows the number of bones overlapping between modules. Statistical significance is shown in color code. Only two intersection were significant about the Bonferroni-corrected *p-value* (yellow to red colors), Ethmoidal + Frontal and Sphenoidal + Occipital, with 9 and 8 nodes overlapping, respectively.

The composition of the two covers resembles that of the anterior and posterior modules reported by other studies using Q modularity optimization (Esteve◻Altava et al., 2013), but the overlapping bones differ from studies using another local optimization approach (Esteve-Altava, 2017a). Here the palatal region connecting the base of the face to the base of the neurocranium is identified as the overlap of the two covers, while in previous reports it was the zygomatic bones who played this role. However, the role of the frontal bone as a bridge between the face and vault is consistently identified using NIMS.

### Modularity of the Human Skull Network with Natural Variation

NIMS returns also four non-redundant node-level modules, here labeled as Wormian, Sphenoidal, Zygomatic Right, and Ethmoidal. The statistical analysis of their intersection returns two statistically significant merges: Ethmoidal + Zygomatic Right and Sphenoidal + Wormian (**Figure 3**). The modules resulting from merging the two pairs of node-level modules are the same than for the ‘type’ human skull network. Likewise, both covers meet the standard definition of module (facial cover: *W* = 160, *p-value* = 4.42e-05; cranial cover: *W* = 146.5, *p-value* = 6.29e-04). Therefore, NIMS returns robust modules even in the presence of natural variation on anatomical networks.

**Figure 3.**
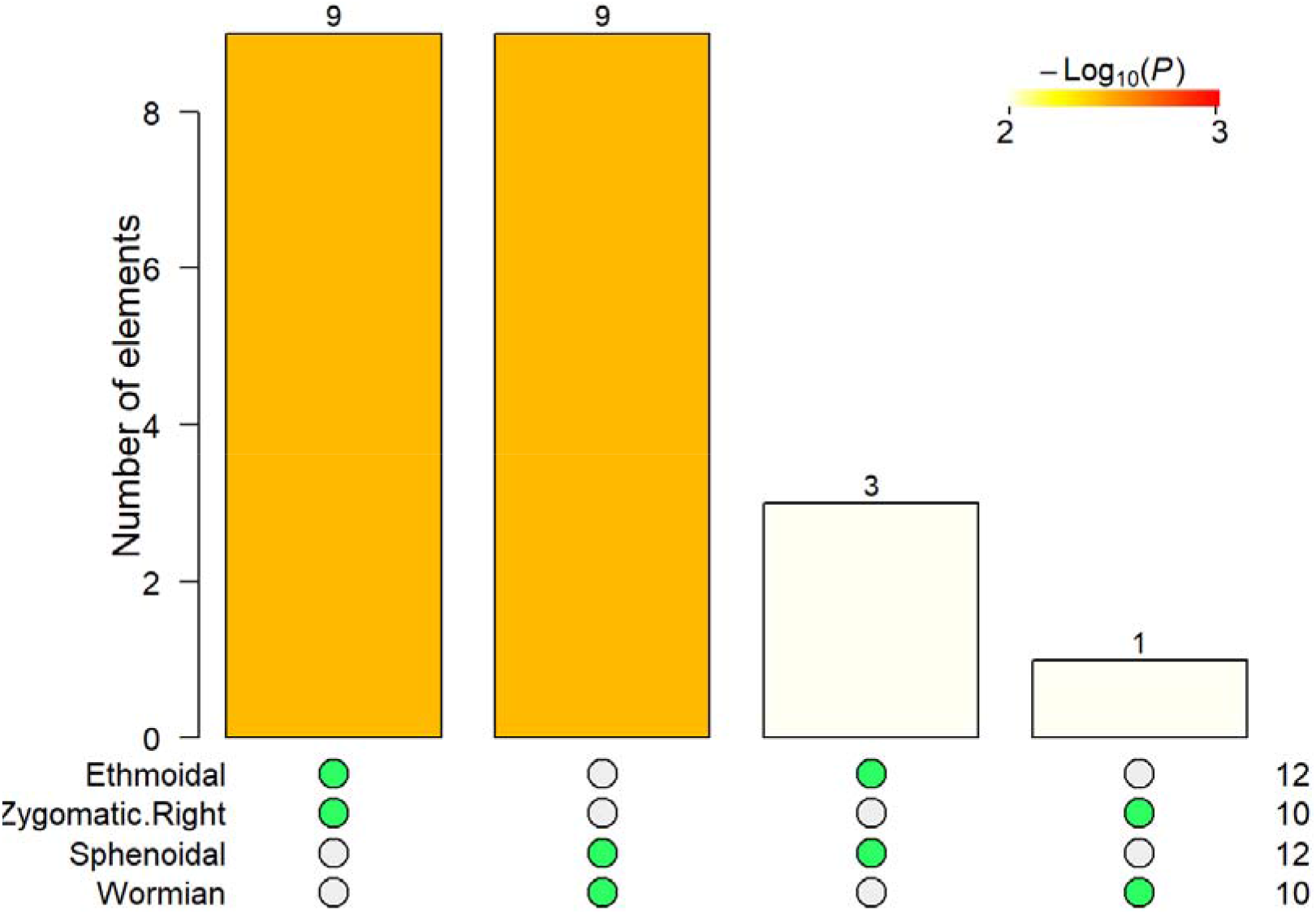
Intersection tests for the four non-redundant node-level modules of the human skull network with variation. See figure 2 for description of the legends.

### Modularity of the Tinamou Skull Network

As expected, NIMS returned one single node-level module grouping the six bones after removing redundancies (all single node-level grouped the same bones). Therefore, no further analysis was required.

### Modularity of the Crocodile Skull Network

NIMS returns seven non-redundant node-level modules, labeled following nodes’ name as R.Postorbital, L.Postorbital, R.Squamosal, Frontal, R.Vomer, L.Vomer, and Pterygoid. The statistical analysis of the intersections for the seven modules returned two significant overlaps (**Figure 4**), which indicate that we can merge these node-level modules. One merging joined R.Vomer + L.Vomer which share 20 of their 21 nodes. Another merging joined R.Postorbital + L.Postorbital + R.Squamosal which share 15 of their 19, 19, and 17 nodes, respectively. Finally, node-level modules Frontal and Pterygoid do not significatively overlap with other modules in a way that improves other merges. Note that a potential merging Pterygoid + R.Postorbital + L.Postorbital + R.Squamosal has not only a lower statistical support (as compared to the merge without Pterygoid) but also reduces the overall size of the overlap from 15 to 8 nodes. Consequently, the merging excluding the Pterygoid is preferred.

**Figure 4.**
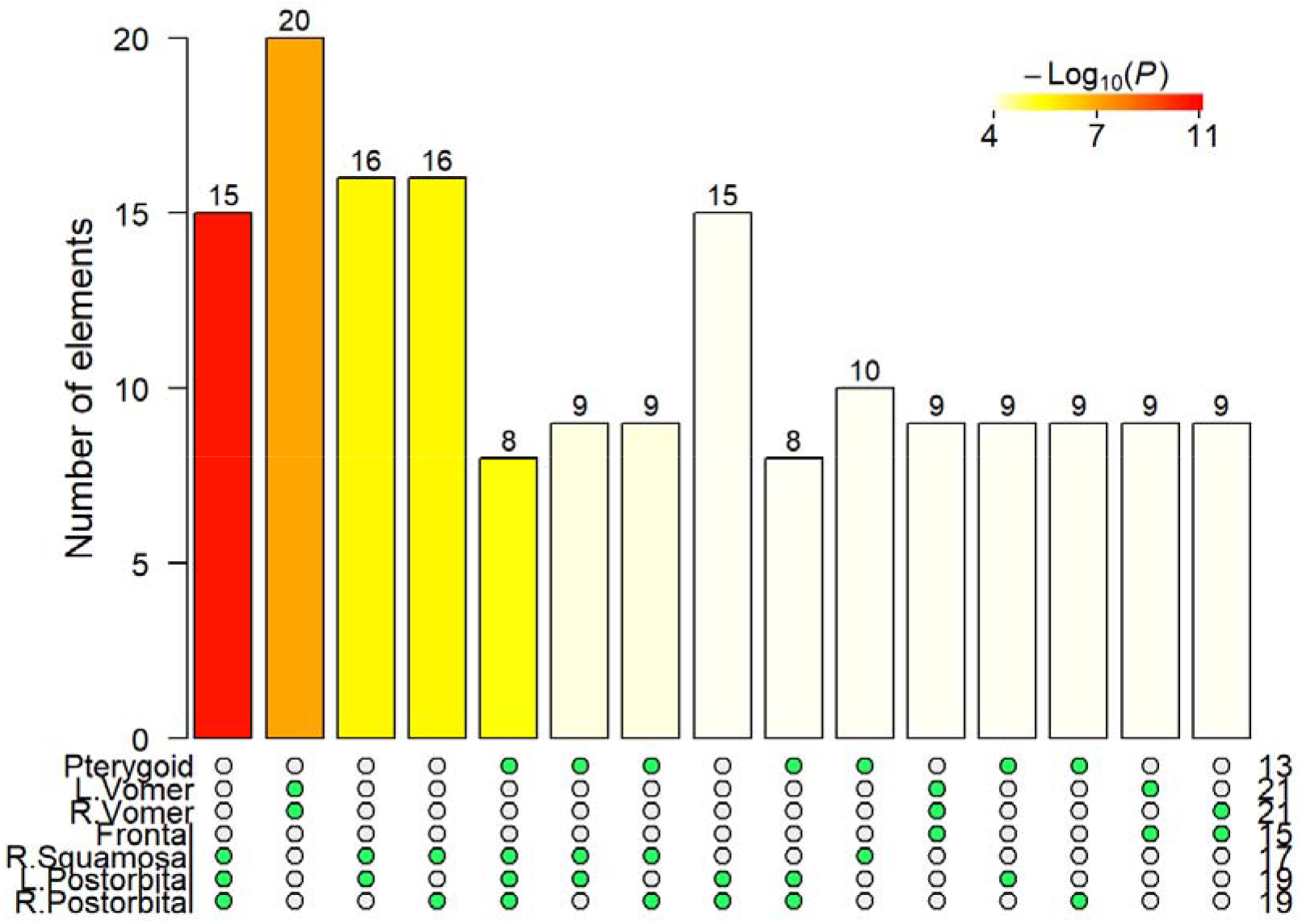
Intersection tests for the seven non-redundant node-level modules of crocodile skull. See figure 2 for description of the legends.

After the two merges, we ended up with four network covers, labeled as frontal, postorbital, pterygoid, and vomer. Frontal module covers the frontal, lacrimals, laterosphenoids, nasals, parietal, postorbitals, prefrontals, squamosals, and supraoccipital. Postorbital module covers the basioccipital, basisphenoid, ectopterygoids, jugals, laterosphenoids, otoccipitals, parietal, postorbitals, prootics, pterygoid, quadrates, quadratojugals, squamosals, and supraoccipital. Pterygoid module covers the basisphenoid, laterosphenoids, palatine, prootics, pterygoid, quadrates, quadratojugals, and vomers. Vomer module covers the ectopterygoids, frontal, jugals, lacrimals, maxillas, nasals, palatine, postorbitals, prefrontals, premaxillas, quadratojugals, and vomers.

The overlap between covers range from two to ten bones (see **Supplementary File 1** for details). Only two covers, vomer and postorbital, show some bones that are exclusive of these modules and do not overlap. Despite having four modules, the skull of the crocodile is highly integrated due to a substantial overlap. It is not surprising that whole-network optimization approaches often return asymmetries since overlapping bones can be placed equally well in different modules. However, the four covers reported here show no asymmetries; additionally, all of them have a bilateral symmetry and group the same bones from the left and right sides. Finally, statistical evaluation of the four covers confirms they meet the standard definition of module (frontal cover: *W* = 191, *p-value* = 3.41e-04; postorbital cover: *W* = 522.5, *p-value* = 4.6e-09; pterygoid cover: *W* = 142.5, *p-value* = 1.37e-03; vomer cover: *W* = 467, *p-value* = 3.84e-08).

## CONCLUSION

NIMS successfully resolved the modular organization of the four skull networks used as example of different types of anatomical networks (**Figure 5**). It found the same modules in ‘type’ networks and in networks including intraspecific variation. It also identified absence of modular organization due to small size and filters out artificial asymmetries. By working under supervision (Step 5), NIMS gives researchers back the control of the modularity analysis to decide the most biologically meaningful output.

**Figure 5.**
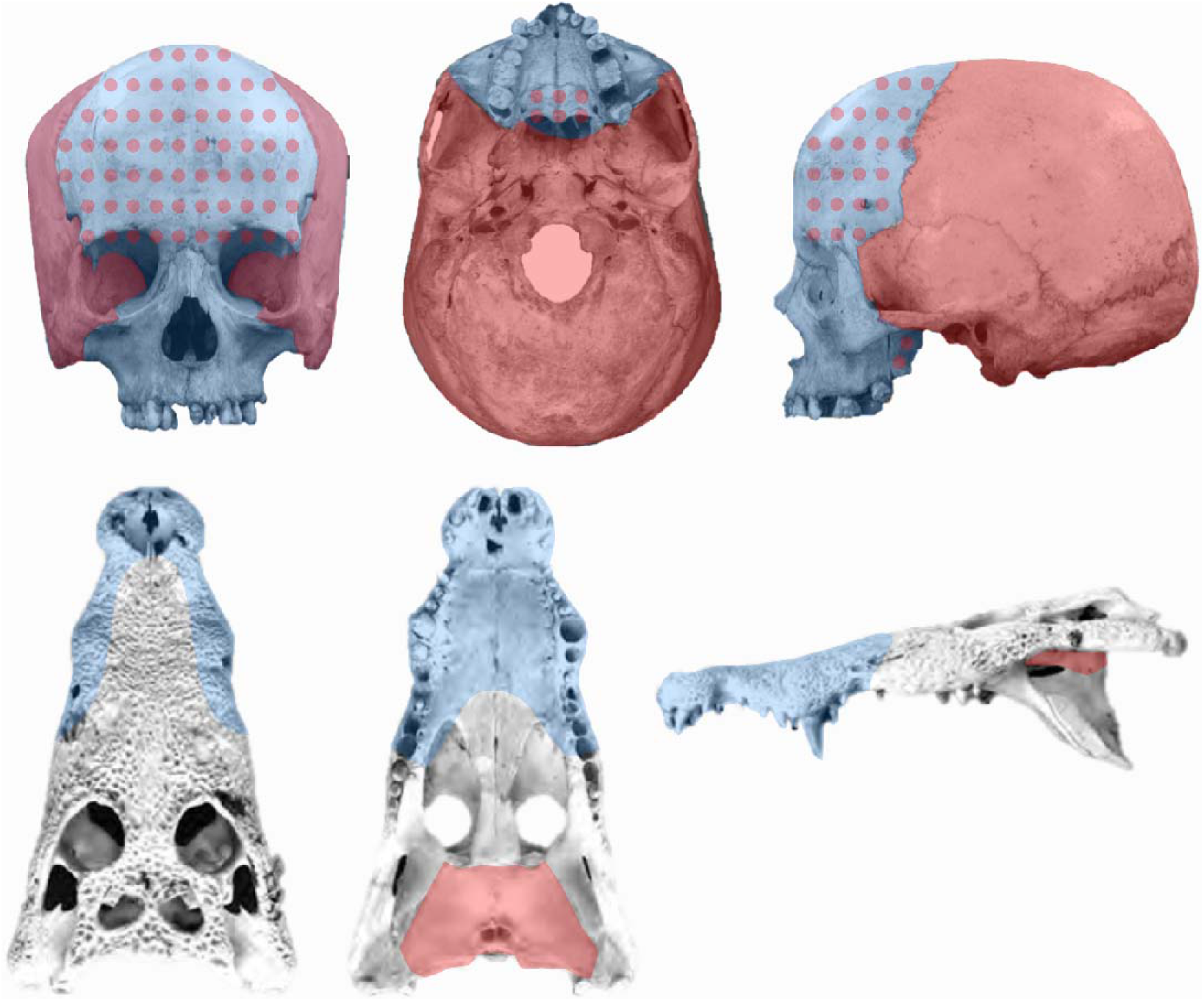
Colored networks by modules. For the human skull network (top), NIMS delimits a posterior cover grouping the cranial vault and base (in red) and an anterior cover delimiting the face (in blue), overlapping in the frontal, palatines, and vomer (red dots on blue background). For the crocodile skull network (bottom), NIMS delimits four covers with a substantial overlap (grey). Only two covers, vomer and postorbital, group bones without overlap (blue and red, respectively). Background image of Homo sapiens adapted from (Takahashi, Yamashita, & Shigehara, 2006) and *Crocodylus moreletii* adapted from (Morgan et al., 2018).

## Acknowledgement

This has received financial support through the Postdoctoral Junior Leader Fellowship Programme from “la Caixa” Banking Foundation (LCF/BQ/LI18/11630002).

## Data and Code Availability

Data and code are included as supplementary materials in Supplementary File 1.

## Authors Contribution

BE-A designed the study, performed the analyses, and wrote the manuscript.

