## Supplementary File 1 for "A Node-based Informed Modularity Strategy to Identify Organizational Modules in Anatomical Networks"

Borja Esteve-Altava

01/07/2020

### Introduction

This is an R Markdown document with the code to reproduce the examples shown in the main text.

#### Load required packages

```
library(igraph)            # network analysis
library(numbers)           # get Bell number
library(SuperExactTest)    # multi-set comparison
```

#### Load anatomical networks

1. Human skull
2. Human skull with variation
3. Tinamou skull
4. Crocodile skull

```
load(url("https://ndownloader.figshare.com/files/23552735"))
```

#### Quick look at the networks

```
for (i in 1:length(networks)){print(networks[i]); print(graph.list[[i]])}
```

```
## [1] "Homo_skull"
## IGRAPH 1f38b7c UN-- 21 64 --
## + attr: name (v/c)
## + edges from 1f38b7c (vertex names):
##  [1] Occipital     --Parietal.Left   Occipital     --Parietal.Right
##  [3] Occipital     --Temporal.Left   Occipital     --Temporal.Right
##  [5] Occipital     --Sphenoidal      Parietal.Left --Parietal.Right
##  [7] Parietal.Left --Temporal.Left   Parietal.Left --Sphenoidal
##  [9] Parietal.Left --Frontal         Parietal.Right--Temporal.Right
## [11] Parietal.Right--Sphenoidal      Parietal.Right--Frontal
## [13] Temporal.Left --Sphenoidal      Temporal.Left --Zygomatic.Left
## [15] Temporal.Right--Sphenoidal      Temporal.Right--Zygomatic.Right
## + ... omitted several edges
## [1] "Homo_skull_variation"
## IGRAPH 1f38deb UN-- 22 66 --
## + attr: name (v/c)
## + edges from 1f38deb (vertex names):
##  [1] Wormian       --Occipital       Wormian       --Parietal.Left
##  [3] Occipital     --Parietal.Left   Occipital     --Parietal.Right
##  [5] Occipital     --Temporal.Left   Occipital     --Temporal.Right
##  [7] Occipital     --Sphenoidal      Parietal.Left --Parietal.Right
##  [9] Parietal.Left --Temporal.Left   Parietal.Left --Sphenoidal
## [11] Parietal.Left --Frontal         Parietal.Right--Temporal.Right
## [13] Parietal.Right--Frontal         Temporal.Left --Sphenoidal
## [15] Temporal.Left --Zygomatic.Left  Temporal.Right--Sphenoidal
## + ... omitted several edges
## [1] "Nothura_skull"
## IGRAPH 1f3905c UN-- 6 9 --
## + attr: name (v/c)
## + edges from 1f3905c (vertex names):
## [1] R.Jugal.Bar--R.Quadrate R.Jugal.Bar--Upper.Beak R.Jugal.Bar--Braincase
## [4] L.Jugal.Bar--L.Quadrate L.Jugal.Bar--Upper.Beak L.Jugal.Bar--Braincase
## [7] R.Quadrate --Braincase  L.Quadrate --Braincase  Upper.Beak --Braincase
## [1] "Crocodylus_skull"
## IGRAPH 1f3953a UN-- 37 103 --
## + attr: name (v/c)
## + edges from 1f3953a (vertex names):
##  [1] R.Premaxilla--L.Premaxilla    R.Premaxilla--R.Maxilla
##  [3] R.Premaxilla--R.Nasal         L.Premaxilla--L.Maxilla
##  [5] L.Premaxilla--L.Nasal         R.Maxilla   --L.Maxilla
##  [7] R.Maxilla   --R.Nasal         R.Maxilla   --R.Lacrimal
##  [9] R.Maxilla   --R.Jugal         R.Maxilla   --R.Vomer
## [11] R.Maxilla   --Palatine        R.Maxilla   --R.Ectopterygoid
## [13] L.Maxilla   --L.Nasal         L.Maxilla   --L.Lacrimal
## [15] L.Maxilla   --L.Jugal         L.Maxilla   --L.Vomer
## + ... omitted several edges
```

### Normal human skull network

```r
# find node-based modules
g<-graph.list[[1]]
mod<-list()
for (i in 1:vcount(g)){
  m<-cluster_spinglass(g,vertex=i)
  mod[[i]]<-V(g)$name[m$community]
}
names(mod)<-V(g)$name

# filter out redundancies
clean_mod<-mod
for (i in 1:(length(clean_mod)-1)){
  j<-i+1
  while (j<=length(clean_mod)){
    if (setequal(clean_mod[[i]],clean_mod[[j]])==TRUE) {clean_mod[[j]]<-NA}
    if (all(is.element(clean_mod[[i]],clean_mod[[j]]))==TRUE) {clean_mod[[i]]<-NA}
    if (all(is.element(clean_mod[[j]],clean_mod[[i]]))==TRUE) {clean_mod[[j]]<-NA}
    j<-j+1
  }
}
clean_mod<-clean_mod[!is.na(clean_mod)]
print(clean_mod)
```

```
## $Occipital
## [1] "Occipital"       "Parietal.Left"   "Parietal.Right"  "Temporal.Left"
## [5] "Temporal.Right"  "Sphenoidal"      "Zygomatic.Right" "Zygomatic.Left"
## [9] "Frontal"
##
## $Sphenoidal
##  [1] "Sphenoidal"      "Occipital"       "Parietal.Left"   "Parietal.Right"
##  [5] "Temporal.Left"   "Temporal.Right"  "Zygomatic.Left"  "Zygomatic.Right"
##  [9] "Palatine.Left"   "Palatine.Right"  "Vomer"
##
## $Frontal
##  [1] "Frontal"            "Ethmoidal"          "Nasal.Left"
##  [4] "Nasal.Right"        "Maxilla.Left"       "Maxilla.Right"
##  [7] "Lacrimal.Left"      "Lacrimal.Right"     "Nasal.Concha.Right"
## [10] "Nasal.Concha.Left"
##
## $Ethmoidal
##  [1] "Ethmoidal"          "Nasal.Left"         "Nasal.Right"
##  [4] "Maxilla.Left"       "Maxilla.Right"      "Lacrimal.Left"
##  [7] "Lacrimal.Right"     "Palatine.Left"      "Palatine.Right"
## [10] "Nasal.Concha.Left"  "Nasal.Concha.Right" "Vomer"
```

```r
# check overlap between node-based modules
res<-supertest(clean_mod,n=vcount(g),degree=c(2:length(clean_mod)))
pcorrect<-(0.05/(bell(length(clean_mod))-length(clean_mod)))
plot(res,sort.by='p-value',degree=c(2:length(clean_mod)),
     Layout='landscape',keep.empty.intersections=FALSE,min.intersection.size=1,
     minMinusLog10PValue=abs(log10(pcorrect)))
```

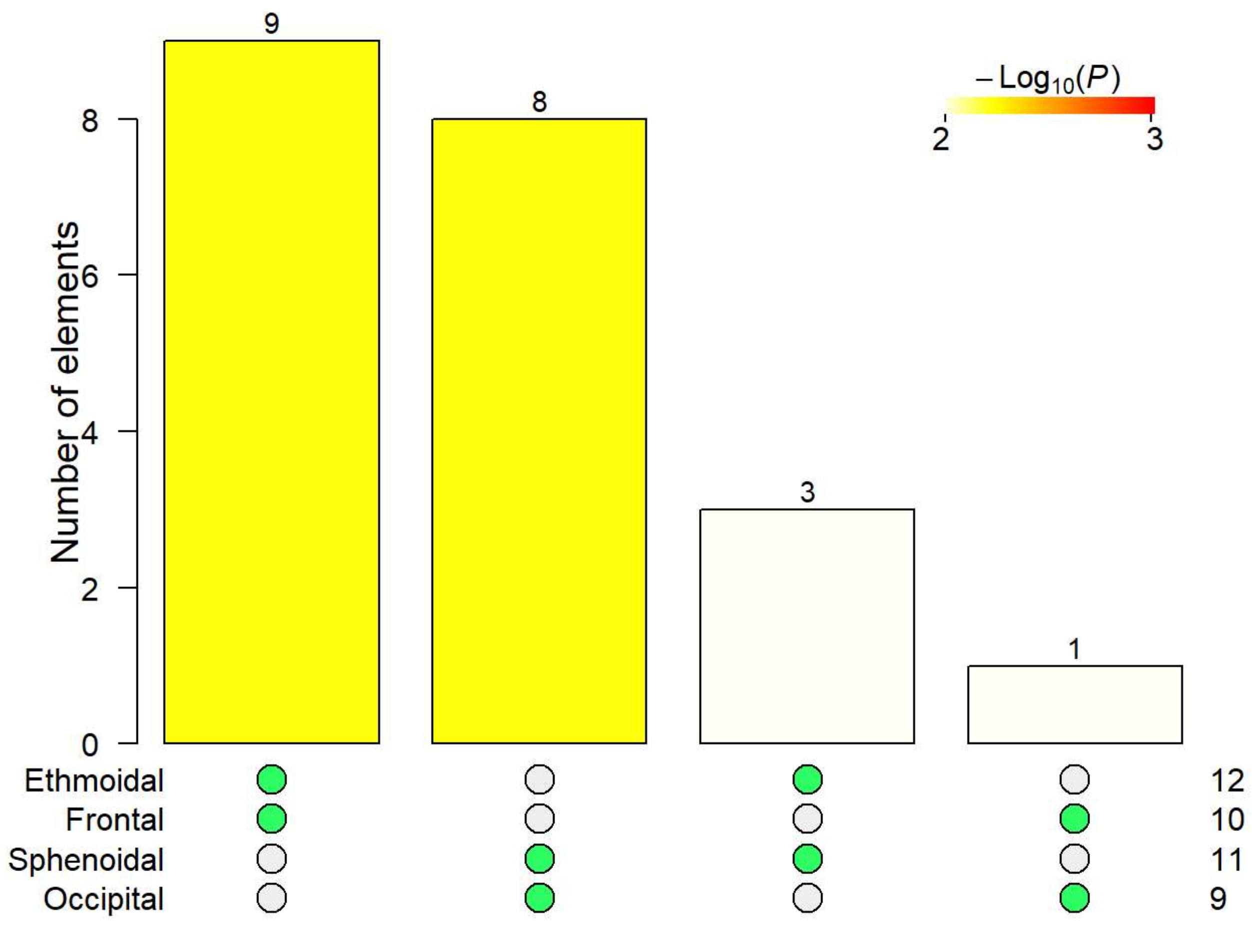


```
# details
facial<-sort(unique(c(clean_mod$Ethmoidal,clean_mod$Frontal)))
cranial<-sort(unique(c(clean_mod$Sphenoidal,clean_mod$Occipital)))
print(facial)
```

```
##  [1] "Ethmoidal"          "Frontal"            "Lacrimal.Left"
##  [4] "Lacrimal.Right"     "Maxilla.Left"       "Maxilla.Right"
##  [7] "Nasal.Concha.Left"  "Nasal.Concha.Right" "Nasal.Left"
## [10] "Nasal.Right"        "Palatine.Left"      "Palatine.Right"
## [13] "Vomer"
```

```
print(cranial)
```

```
##  [1] "Frontal"        "Occipital"      "Palatine.Left"  "Palatine.Right"
##  [5] "Parietal.Left"  "Parietal.Right" "Sphenoidal"     "Temporal.Left"
##  [9] "Temporal.Right" "Vomer"          "Zygomatic.Left" "Zygomatic.Right"
```

```
print(intersect(facial,cranial))
```

```
## [1] "Frontal"        "Palatine.Left"  "Palatine.Right" "Vomer"
```

```
# testing
vids<-c(1:vcount(g))[is.element(V(g)$name,facial)]
in.links<-degree(induced_subgraph(g,vids))
out.links<-degree(g,vids)-in.links
res<-wilcox.test(in.links, out.links, alternative="greater")
print("Facial module")
```

```
## [1] "Facial module"
```

```
print(res)
```

```
## 
##  Wilcoxon rank sum test with continuity correction
## 
## data:  in.links and out.links
## W = 161.5, p-value = 3.124e-05
## alternative hypothesis: true location shift is greater than 0
```

```
vids<-c(1:vcount(g))[is.element(V(g)$name,cranial)]
in.links<-degree(induced_subgraph(g,vids))
out.links<-degree(g,vids)-in.links
res<-wilcox.test(in.links, out.links, alternative="greater")
print("Cranial module")
```

```
## [1] "Cranial module"
```

```
print(res)
```

```
## 
##  Wilcoxon rank sum test with continuity correction
## 
## data:  in.links and out.links
## W = 125.5, p-value = 0.0008381
## alternative hypothesis: true location shift is greater than 0
```

### Human skull network with variability

```
# find node-based modules
g<-graph.list[[2]]
mod<-list()
for (i in 1:vcount(g)){
  m<-cluster_spinglass(g,vertex=i)
  mod[[i]]<-V(g)$name[m$community]
}
names(mod)<-V(g)$name

# filter out redundancies
clean_mod<-mod
for (i in 1:(length(clean_mod)-1)){
  j<-i+1
  while (j<=length(clean_mod)){
    if (setequal(clean_mod[[i]],clean_mod[[j]])==TRUE) {clean_mod[[j]]<-NA}
    if (all(is.element(clean_mod[[i]],clean_mod[[j]]))==TRUE) {clean_mod[[i]]<-NA}
    if (all(is.element(clean_mod[[j]],clean_mod[[i]]))==TRUE) {clean_mod[[j]]<-NA}
    j<-j+1
  }
}
clean_mod<-clean_mod[!is.na(clean_mod)]
print(clean_mod)
```

```
## $Wormian
##  [1] "Wormian"         "Occipital"       "Parietal.Left"   "Temporal.Left"
##  [5] "Parietal.Right"  "Temporal.Right"  "Sphenoidal"      "Zygomatic.Right"
##  [9] "Zygomatic.Left"  "Frontal"
##
## $Sphenoidal
##  [1] "Sphenoidal"      "Occipital"       "Parietal.Left"   "Temporal.Left"
##  [5] "Temporal.Right"  "Zygomatic.Left"  "Zygomatic.Right" "Palatine.Left"
##  [9] "Palatine.Right"  "Vomer"           "Parietal.Right"  "Wormian"
##
## $Zygomatic.Right
##  [1] "Frontal"            "Maxilla.Right"      "Lacrimal.Right"
##  [4] "Nasal.Concha.Right" "Nasal.Right"        "Nasal.Left"
##  [7] "Ethmoidal"          "Lacrimal.Left"      "Maxilla.Left"
## [10] "Nasal.Concha.Left"
##
## $Ethmoidal
##  [1] "Ethmoidal"          "Nasal.Left"         "Nasal.Right"
##  [4] "Maxilla.Left"       "Maxilla.Right"      "Lacrimal.Left"
##  [7] "Lacrimal.Right"     "Palatine.Left"      "Palatine.Right"
## [10] "Nasal.Concha.Left"  "Nasal.Concha.Right" "Vomer"
```

```
# check overlap between node-based modules
res<-supertest(clean_mod,n=vcount(g),degree=c(2:length(clean_mod)))
pcorrect<-(0.05/(bell(length(clean_mod))-length(clean_mod)))
plot(res,sort.by='p-value',degree=c(2:length(clean_mod)),
     Layout='landscape',keep.empty.intersections=FALSE,min.intersection.size=1,
     minMinusLog10PValue=abs(log10(pcorrect)))
```

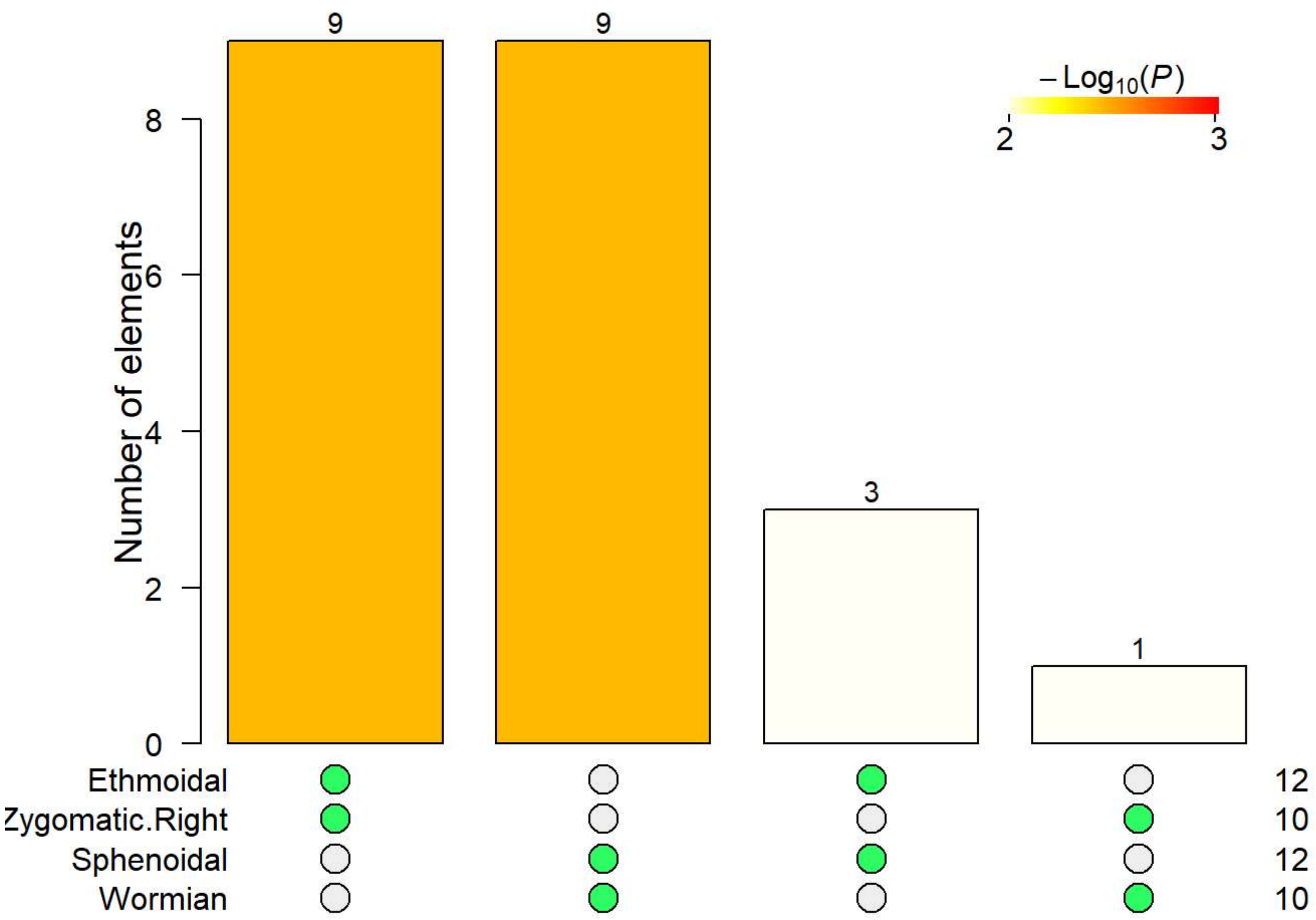


```
# details
facial<-sort(unique(c(clean_mod$Ethmoidal,clean_mod$Zygomatic.Right)))
cranial<-sort(unique(c(clean_mod$Sphenoidal,clean_mod$Wormian)))
print(facial)
```

```
##  [1] "Ethmoidal"          "Frontal"            "Lacrimal.Left"
##  [4] "Lacrimal.Right"     "Maxilla.Left"       "Maxilla.Right"
##  [7] "Nasal.Concha.Left"  "Nasal.Concha.Right" "Nasal.Left"
## [10] "Nasal.Right"        "Palatine.Left"      "Palatine.Right"
## [13] "Vomer"
```

```
print(cranial)
```

```
##  [1] "Frontal"         "Occipital"       "Palatine.Left"   "Palatine.Right"
##  [5] "Parietal.Left"   "Parietal.Right"  "Sphenoidal"      "Temporal.Left"
##  [9] "Temporal.Right"  "Vomer"           "Wormian"         "Zygomatic.Left"
## [13] "Zygomatic.Right"
```

```
print(intersect(facial,cranial))
```

```
## [1] "Frontal"        "Palatine.Left"  "Palatine.Right" "Vomer"
```

```r
# testing
vids<-c(1:vcount(g))[is.element(V(g)$name,facial)]
in.links<-degree(induced_subgraph(g,vids))
out.links<-degree(g,vids)-in.links
res<-wilcox.test(in.links, out.links, alternative="greater")
print("Facial module")
```

```
## [1] "Facial module"
```

```r
print(res)
```

```
##
##  Wilcoxon rank sum test with continuity correction
##
## data:  in.links and out.links
## W = 160, p-value = 4.417e-05
## alternative hypothesis: true location shift is greater than 0
```

```r
vids<-c(1:vcount(g))[is.element(V(g)$name,cranial)]
in.links<-degree(induced_subgraph(g,vids))
out.links<-degree(g,vids)-in.links
res<-wilcox.test(in.links, out.links, alternative="greater")
print("Cranial module")
```

```
## [1] "Cranial module"
```

```r
print(res)
```

```
##
##  Wilcoxon rank sum test with continuity correction
##
## data:  in.links and out.links
## W = 146.5, p-value = 0.0006294
## alternative hypothesis: true location shift is greater than 0
```

### Tinamou skull network

```
# find node-based modules
g<-graph.list[[3]]
mod<-list()
for (i in 1:vcount(g)){
  m<-cluster_spinglass(g,vertex=i)
  mod[[i]]<-V(g)$name[m$community]
}
names(mod)<-V(g)$name

# filter out redundancies
clean_mod<-mod
for (i in 1:(length(clean_mod)-1)){
  j<-i+1
  while (j<=length(clean_mod)){
    if (setequal(clean_mod[[i]],clean_mod[[j]])==TRUE) {clean_mod[[j]]<-NA}
    if (all(is.element(clean_mod[[i]],clean_mod[[j]]))==TRUE) {clean_mod[[i]]<-NA}
    if (all(is.element(clean_mod[[j]],clean_mod[[i]]))==TRUE) {clean_mod[[j]]<-NA}
    j<-j+1
  }
}
clean_mod<-clean_mod[!is.na(clean_mod)]
print(clean_mod)
```

```
## $Braincase
## [1] "Braincase"   "R.Jugal.Bar" "L.Jugal.Bar" "R.Quadrate"  "L.Quadrate"
## [6] "Upper.Beak"
```

### Crocodile skull network

```
# find node-based modules
g<-graph.list[[4]]
mod<-list()
for (i in 1:vcount(g)){
  m<-cluster_spinglass(g,vertex=i)
  mod[[i]]<-V(g)$name[m$community]
}
names(mod)<-V(g)$name

# filter out redundancies
clean_mod<-mod
for (i in 1:(length(clean_mod)-1)){
  j<-i+1
  while (j<=length(clean_mod)){
    if (setequal(clean_mod[[i]],clean_mod[[j]])==TRUE) {clean_mod[[j]]<-NA}
    if (all(is.element(clean_mod[[i]],clean_mod[[j]]))==TRUE) {clean_mod[[i]]<-NA}
    if (all(is.element(clean_mod[[j]],clean_mod[[i]]))==TRUE) {clean_mod[[j]]<-NA}
    j<-j+1
  }
}
clean_mod<-clean_mod[!is.na(clean_mod)]
print(clean_mod)
```

```
## $R.Postorbital
##  [1] "R.Postorbital"    "R.Jugal"          "R.Squamosal"      "Parietal"
##  [5] "R.Ectopterygoid"  "L.Squamosal"      "Supraoccipital"   "R.Quadratojugal"
##  [9] "R.Quadrate"       "R.Otoccipital"    "L.Otoccipital"    "Basioccipital"
## [13] "R.Laterosphenoid" "Basisphenoid"     "R.Prootic"        "L.Laterosphenoid"
## [17] "L.Quadrate"       "L.Prootic"        "Pterygoid"
##
## $L.Postorbital
##  [1] "L.Postorbital"    "L.Jugal"          "L.Squamosal"      "Parietal"
##  [5] "L.Ectopterygoid"  "R.Squamosal"      "Supraoccipital"   "L.Quadratojugal"
##  [9] "L.Quadrate"       "L.Otoccipital"    "R.Otoccipital"    "Basioccipital"
## [13] "L.Laterosphenoid" "Basisphenoid"     "L.Prootic"        "R.Laterosphenoid"
## [17] "R.Quadrate"       "R.Prootic"        "Pterygoid"
##
## $R.Squamosal
##  [1] "R.Squamosal"      "Parietal"         "Supraoccipital"   "R.Otoccipital"
##  [5] "R.Quadrate"       "L.Squamosal"      "L.Otoccipital"    "Basioccipital"
##  [9] "Basisphenoid"     "L.Quadrate"       "L.Prootic"        "L.Laterosphenoid"
## [13] "R.Laterosphenoid" "R.Prootic"        "Pterygoid"        "L.Quadratojugal"
## [17] "R.Quadratojugal"
##
## $Frontal
##  [1] "Frontal"          "R.Nasal"          "L.Nasal"          "R.Prefrontal"
##  [5] "L.Prefrontal"     "R.Postorbital"    "L.Postorbital"    "Parietal"
##  [9] "R.Laterosphenoid" "L.Laterosphenoid" "L.Lacrimal"       "R.Lacrimal"
## [13] "L.Squamosal"      "Supraoccipital"   "R.Squamosal"
##
## $R.Vomer
##  [1] "R.Vomer"         "R.Maxilla"       "L.Vomer"         "Palatine"
##  [5] "L.Maxilla"       "L.Ectopterygoid" "R.Ectopterygoid" "L.Jugal"
##  [9] "L.Lacrimal"      "L.Prefrontal"    "L.Nasal"         "L.Premaxilla"
## [13] "R.Premaxilla"    "R.Nasal"         "R.Prefrontal"    "R.Lacrimal"
## [17] "R.Jugal"         "L.Quadratojugal" "Frontal"         "R.Postorbital"
## [21] "L.Postorbital"
##
## $L.Vomer
##  [1] "L.Vomer"         "L.Maxilla"       "R.Vomer"         "Palatine"
##  [5] "R.Maxilla"       "R.Ectopterygoid" "L.Ectopterygoid" "R.Jugal"
##  [9] "R.Lacrimal"      "R.Prefrontal"    "R.Nasal"         "R.Premaxilla"
## [13] "L.Premaxilla"    "L.Nasal"         "L.Prefrontal"    "L.Lacrimal"
## [17] "L.Jugal"         "R.Quadratojugal" "Frontal"         "L.Postorbital"
## [21] "R.Postorbital"
##
## $Pterygoid
##  [1] "Pterygoid"        "R.Prootic"        "L.Prootic"        "Basisphenoid"
##  [5] "R.Laterosphenoid" "L.Laterosphenoid" "R.Vomer"          "L.Vomer"
##  [9] "Palatine"         "R.Quadrate"       "L.Quadrate"       "L.Quadratojugal"
## [13] "R.Quadratojugal"
```

```
# check overlap between node-based modules
res<-supertest(clean_mod,n=vcount(g),degree=c(2:length(clean_mod)))
pcorrect<-(0.05/(bell(length(clean_mod))-length(clean_mod)))
plot(res,sort.by='p-value',degree=c(2:length(clean_mod)),
     Layout='landscape',keep.empty.intersections=FALSE,min.intersection.size=8,
     minMinusLog10PValue=abs(log10(pcorrect)))
```

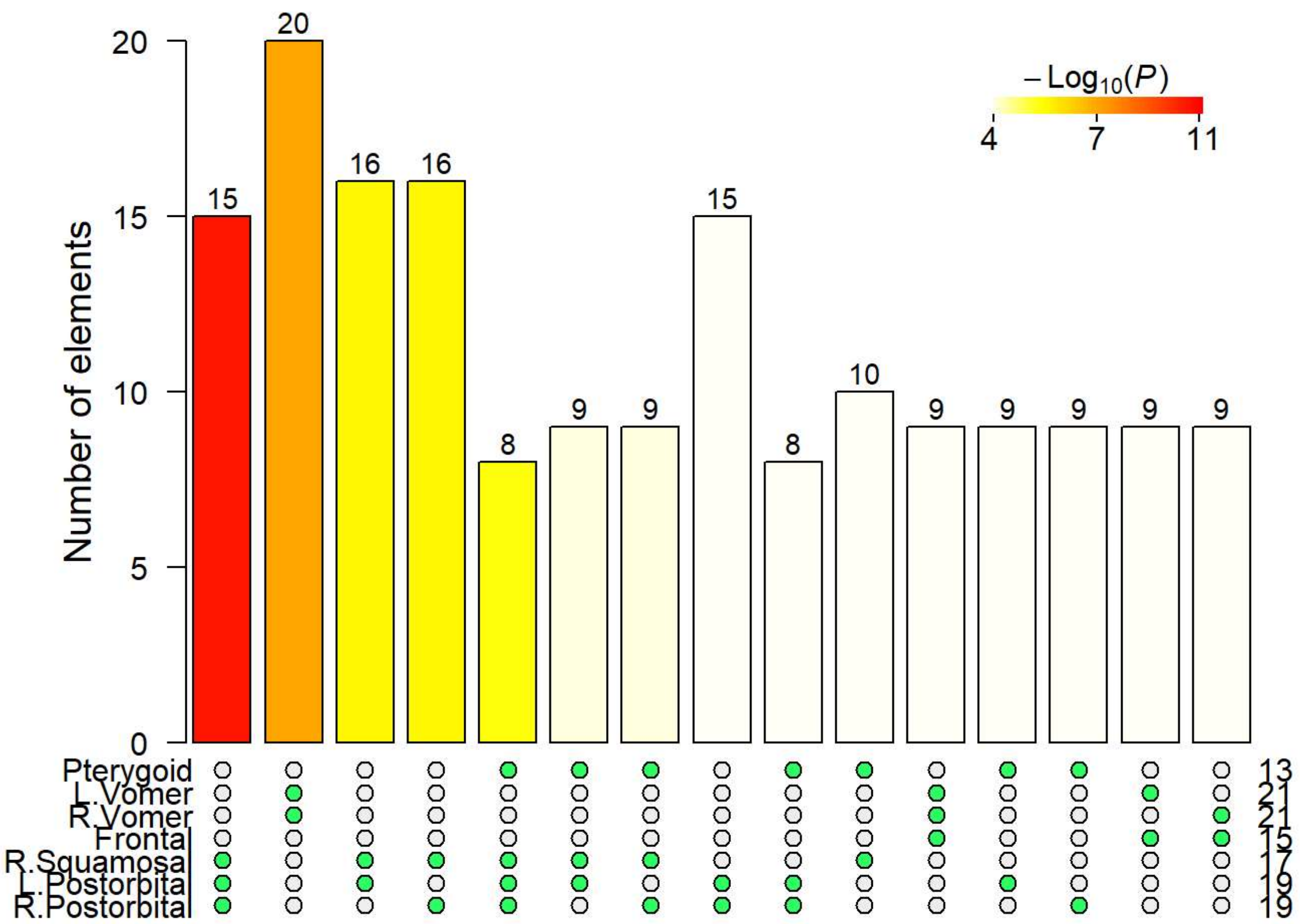


```
# details
pterygoid<-sort(unique(c(clean_mod$Pterygoid)))
frontal<-sort(unique(c(clean_mod$Frontal)))
postorbital<-sort(unique(c(clean_mod$R.Squamosal,clean_mod$R.Postorbital,clean_mod$L.Postorbi
tal)))
vomer<-sort(unique(c(clean_mod$R.Vomer,clean_mod$L.Vomer)))
print(pterygoid)
```

```
##  [1] "Basisphenoid"     "L.Laterosphenoid" "L.Prootic"        "L.Quadrate"
##  [5] "L.Quadratojugal"  "L.Vomer"          "Palatine"         "Pterygoid"
##  [9] "R.Laterosphenoid" "R.Prootic"        "R.Quadrate"       "R.Quadratojugal"
## [13] "R.Vomer"
```

```
print(frontal)
```

```
##  [1] "Frontal"          "L.Lacrimal"       "L.Laterosphenoid" "L.Nasal"
##  [5] "L.Postorbital"    "L.Prefrontal"     "L.Squamosal"      "Parietal"
##  [9] "R.Lacrimal"       "R.Laterosphenoid" "R.Nasal"          "R.Postorbital"
## [13] "R.Prefrontal"     "R.Squamosal"      "Supraocciptal"
```

```
print(postorbital)
```

```
##  [1] "Basioccipital"    "Basisphenoid"     "L.Ectopterygoid"  "L.Jugal"
##  [5] "L.Laterosphenoid" "L.Otoccipital"    "L.Postorbital"    "L.Prootic"
##  [9] "L.Quadrate"       "L.Quadratojugal"  "L.Squamosal"      "Parietal"
## [13] "Pterygoid"        "R.Ectopterygoid"  "R.Jugal"          "R.Laterosphenoid"
## [17] "R.Otoccipital"    "R.Postorbital"    "R.Prootic"        "R.Quadrate"
## [21] "R.Quadratojugal"  "R.Squamosal"      "Supraocciptal"
```

```
print(vomer)
```

```
##  [1] "Frontal"         "L.Ectopterygoid" "L.Jugal"         "L.Lacrimal"
##  [5] "L.Maxilla"       "L.Nasal"         "L.Postorbital"   "L.Prefrontal"
##  [9] "L.Premaxilla"    "L.Quadratojugal" "L.Vomer"         "Palatine"
## [13] "R.Ectopterygoid" "R.Jugal"         "R.Lacrimal"      "R.Maxilla"
## [17] "R.Nasal"         "R.Postorbital"   "R.Prefrontal"    "R.Premaxilla"
## [21] "R.Quadratojugal" "R.Vomer"
```

```
intersect(pterygoid,frontal)
```

```
## [1] "L.Laterosphenoid" "R.Laterosphenoid"
```

```
intersect(pterygoid,postorbital)
```

```
##  [1] "Basisphenoid"     "L.Laterosphenoid" "L.Prootic"        "L.Quadrate"
##  [5] "L.Quadratojugal"  "Pterygoid"        "R.Laterosphenoid" "R.Prootic"
##  [9] "R.Quadrate"       "R.Quadratojugal"
```

```
intersect(pterygoid,vomer)
```

```
## [1] "L.Quadratojugal" "L.Vomer"         "Palatine"        "R.Quadratojugal"
## [5] "R.Vomer"
```

```
intersect(frontal,postorbital)
```

```
## [1] "L.Laterosphenoid" "L.Postorbital"    "L.Squamosal"      "Parietal"
## [5] "R.Laterosphenoid" "R.Postorbital"    "R.Squamosal"      "Supraocciptal"
```

```
intersect(frontal,vomer)
```

```
## [1] "Frontal"       "L.Lacrimal"    "L.Nasal"       "L.Postorbital"
## [5] "L.Prefrontal"  "R.Lacrimal"    "R.Nasal"       "R.Postorbital"
## [9] "R.Prefrontal"
```

```
intersect(postorbital,vomer)
```

```
## [1] "L.Ectopterygoid" "L.Jugal"         "L.Postorbital"   "L.Quadratojugal"
## [5] "R.Ectopterygoid" "R.Jugal"         "R.Postorbital"   "R.Quadratojugal"
```

```
# testing
vids<-c(1:vcount(g))[is.element(V(g)$name,frontal)]
in.links<-degree(induced_subgraph(g,vids))
out.links<-degree(g,vids)-in.links
res<-wilcox.test(in.links, out.links, alternative="greater")
print("Frontal module")
```

```
## [1] "Frontal module"
```

```
print(res)
```

```
##
##  Wilcoxon rank sum test with continuity correction
##
## data:  in.links and out.links
## W = 191, p-value = 0.000341
## alternative hypothesis: true location shift is greater than 0
```

```
vids<-c(1:vcount(g))[is.element(V(g)$name,postorbital)]
in.links<-degree(induced_subgraph(g,vids))
out.links<-degree(g,vids)-in.links
res<-wilcox.test(in.links, out.links, alternative="greater")
print("Postorbital module")
```

```
## [1] "Postorbital module"
```

```
print(res)
```

```
##
##  Wilcoxon rank sum test with continuity correction
##
## data:  in.links and out.links
## W = 522.5, p-value = 4.604e-09
## alternative hypothesis: true location shift is greater than 0
```

```
vids<-c(1:vcount(g))[is.element(V(g)$name,pterygoid)]
in.links<-degree(induced_subgraph(g,vids))
out.links<-degree(g,vids)-in.links
res<-wilcox.test(in.links, out.links, alternative="greater")
print("Pterygoid module")
```

```
## [1] "Pterygoid module"
```

```
print(res)
```

```
##
##  Wilcoxon rank sum test with continuity correction
##
## data:  in.links and out.links
## W = 142.5, p-value = 0.001374
## alternative hypothesis: true location shift is greater than 0
```

```
vids<-c(1:vcount(g))[is.element(V(g)$name,vomer)]
in.links<-degree(induced_subgraph(g,vids))
out.links<-degree(g,vids)-in.links
res<-wilcox.test(in.links, out.links, alternative="greater")
print("Vomer module")
```

```
## [1] "Vomer module"
```

```
print(res)
```

```
##
##  Wilcoxon rank sum test with continuity correction
##
## data:  in.links and out.links
## W = 467, p-value = 3.838e-08
## alternative hypothesis: true location shift is greater than 0
```

### Zachary’s karate club network

```
library(igraphdata)
data(karate,package="igraphdata")
# find node-based modules
g<-karate
mod<-list()
for (i in 1:vcount(g)){
  m<-cluster_spinglass(g,vertex=i)
  mod[[i]]<-V(g)$name[m$community]
}
names(mod)<-V(g)$name

# filter out redundancies
clean_mod<-mod
for (i in 1:(length(clean_mod)-1)){
  j<-i+1
  while (j<=length(clean_mod)){
    if (setequal(clean_mod[[i]],clean_mod[[j]])==TRUE) {clean_mod[[j]]<-NA}
    if (all(is.element(clean_mod[[i]],clean_mod[[j]]))==TRUE) {clean_mod[[i]]<-NA}
    if (all(is.element(clean_mod[[j]],clean_mod[[i]]))==TRUE) {clean_mod[[j]]<-NA}
    j<-j+1
  }
}
clean_mod<-clean_mod[!is.na(clean_mod)]
print(clean_mod)
```

```
## $`Mr Hi`
##  [1] "Mr Hi"     "Actor 2"  "Actor 3"  "Actor 4"  "Actor 5"  "Actor 6"
##  [7] "Actor 7"  "Actor 8"  "Actor 11" "Actor 12" "Actor 13" "Actor 14"
## [13] "Actor 18" "Actor 20" "Actor 22" "Actor 17"
##
## $`Actor 2`
##  [1] "Actor 2"  "Mr Hi"     "Actor 3"  "Actor 4"  "Actor 8"  "Actor 14"
##  [7] "Actor 18" "Actor 20" "Actor 22" "Actor 31" "Actor 9"  "Actor 13"
## [13] "Actor 12"
##
## $`Actor 9`
##  [1] "Actor 9"  "Actor 31" "Actor 33" "John A"   "Actor 16" "Actor 23"
##  [7] "Actor 15" "Actor 21" "Actor 24" "Actor 30" "Actor 27" "Actor 28"
## [13] "Actor 19" "Actor 10" "Actor 25" "Actor 26" "Actor 32" "Actor 29"
```

```
# check overlap between node-based modules
res<-supertest(clean_mod,n=vcount(g),degree=c(2:length(clean_mod)))
pcorrect<-(0.05/(bell(length(clean_mod))-length(clean_mod)))
plot(res,sort.by='p-value',degree=c(2:length(clean_mod)),
     Layout='landscape',keep.empty.intersections=FALSE,min.intersection.size=1,
     minMinusLog10PValue=abs(log10(pcorrect)))
```

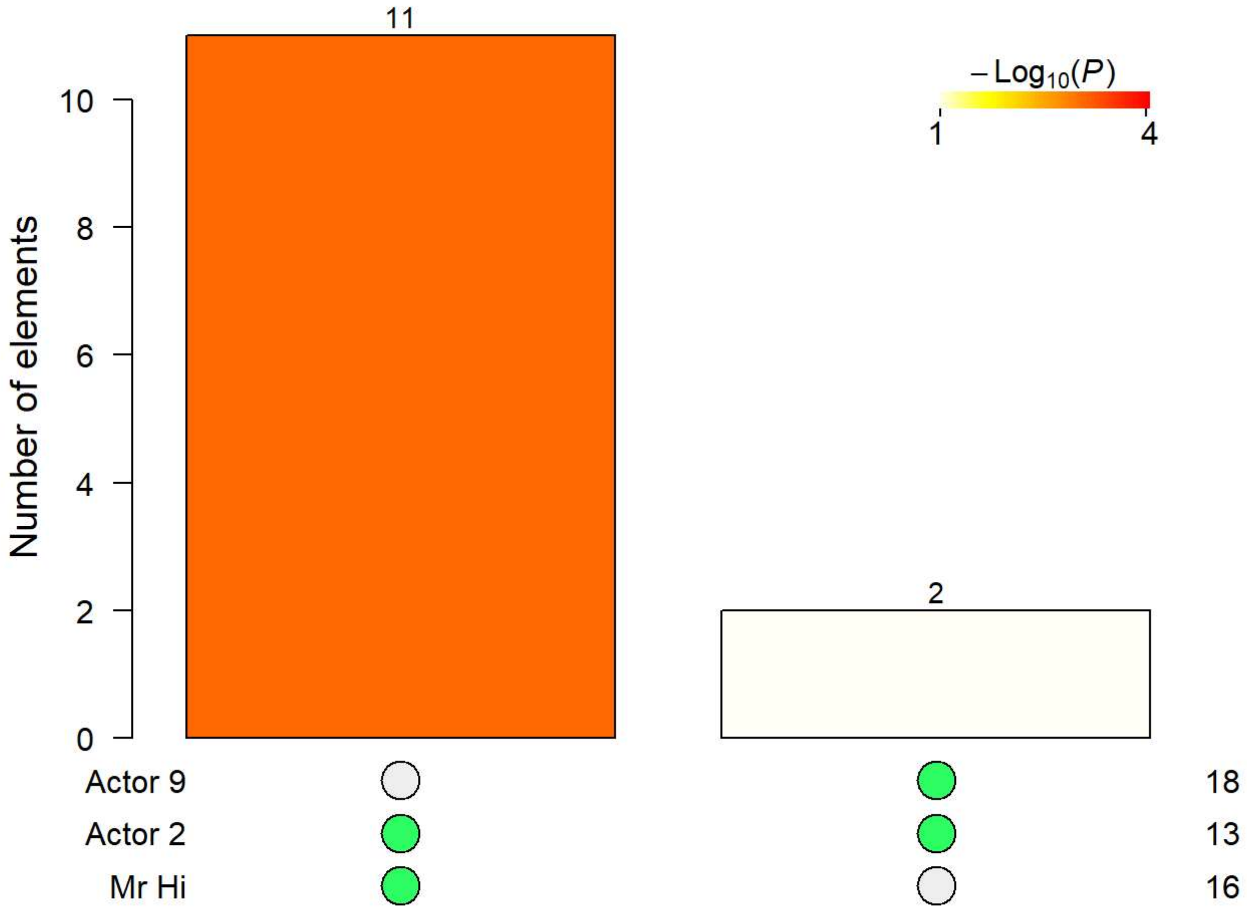


```
# details
moduleA<-sort(unique(c(clean_mod$`Actor 9`)))
moduleB<-sort(unique(c(clean_mod$`Actor 2`,clean_mod$`Mr Hi`)))
print(moduleA)
```

```
##  [1] "Actor 10" "Actor 15" "Actor 16" "Actor 19" "Actor 21" "Actor 23"
##  [7] "Actor 24" "Actor 25" "Actor 26" "Actor 27" "Actor 28" "Actor 29"
## [13] "Actor 30" "Actor 31" "Actor 32" "Actor 33" "Actor 9"  "John A"
```

```
print(moduleB)
```

```
##  [1] "Actor 11" "Actor 12" "Actor 13" "Actor 14" "Actor 17" "Actor 18"
##  [7] "Actor 2"  "Actor 20" "Actor 22" "Actor 3"  "Actor 31" "Actor 4"
## [13] "Actor 5"  "Actor 6"  "Actor 7"  "Actor 8"  "Actor 9"  "Mr Hi"
```

```
print(intersect(moduleA,moduleB))
```

```
## [1] "Actor 31" "Actor 9"
```

```
# compare with the actual groups
mem<-V(g)$Faction; names(mem)<-V(g)$name
print(mem[is.element(V(g)$name,moduleB)])
```

```
##    Mr Hi  Actor 2  Actor 3  Actor 4  Actor 5  Actor 6  Actor 7  Actor 8
##        1        1        1        1        1        1        1        1
##  Actor 9 Actor 11 Actor 12 Actor 13 Actor 14 Actor 17 Actor 18 Actor 20
##        2        1        1        1        1        1        1        1
## Actor 22 Actor 31
##        1        2
```

```
print(mem[is.element(V(g)$name,moduleA)])
```

```
##  Actor 9 Actor 10 Actor 15 Actor 16 Actor 19 Actor 21 Actor 23 Actor 24
##        2        2        2        2        2        2        2        2
## Actor 25 Actor 26 Actor 27 Actor 28 Actor 29 Actor 30 Actor 31 Actor 32
##        2        2        2        2        2        2        2        2
## Actor 33   John A
##        2        2
```

### Session information

```
print(sessionInfo())
```

```
## R version 3.6.2 (2019-12-12)
## Platform: x86_64-w64-mingw32/x64 (64-bit)
## Running under: Windows 10 x64 (build 18363)
##
## Matrix products: default
##
## locale:
## [1] LC_COLLATE=Spanish_Spain.1252  LC_CTYPE=Spanish_Spain.1252
## [3] LC_MONETARY=Spanish_Spain.1252 LC_NUMERIC=C
## [5] LC_TIME=Spanish_Spain.1252
##
## attached base packages:
## [1] grid      stats     graphics  grDevices utils     datasets  methods
## [8] base
##
## other attached packages:
## [1] igraphdata_1.0.1     SuperExactTest_1.0.7 numbers_0.7-5
## [4] igraph_1.2.4.2
##
## loaded via a namespace (and not attached):
##  [1] Rcpp_1.0.3      digest_0.6.23   magrittr_1.5    evaluate_0.14
##  [5] rlang_0.4.6     stringi_1.4.4   rmarkdown_2.1   tools_3.6.2
##  [9] stringr_1.4.0   xfun_0.13       yaml_2.2.1      compiler_3.6.2
## [13] pkgconfig_2.0.3 htmltools_0.4.0 knitr_1.28
```
